# A core UPS molecular complement implicates unique endocytic compartments at the parasite-host interface in *Giardia lamblia*

**DOI:** 10.1101/2022.07.13.499947

**Authors:** Erina A. Balmer, Corina D. Wirdnam, Carmen Faso

**Affiliations:** Institute of Cell Biology, University of Bern, Bern, Switzerland; Multidisciplinary Center for Infectious Diseases, Vetsuisse Faculty, University of Bern, Bern, Switzerland; Graduate School for Cellular and Biomedical Sciences, University of Bern, Bern, Switzerland

**Keywords:** *Giardia lamblia*, virulence, unconventional protein secretion, interactome, peripheral endocytic compartments

## Abstract

Unconventional protein secretion (UPS) plays important roles in processes for the survival of the cell and whole organisms. In contrast to canonical secretory routes, UPS does not generally require secretory signal sequences and often bypasses secretory compartments such as the ER and the Golgi apparatus.

*Giardia lamblia* is a protozoan parasite of global medical importance and reduced subcellular complexity known to release several proteins, some of them virulence factors, without canonical secretory signals, thus implicating UPS at the parasite-host interface. No dedicated machinery nor mechanism(s) for UPS in Giardia are currently known, although speculations on unique endocytic Giardia compartments called PV/PECs have been put forth.

To begin to address the question of whether PV/PECs are implicated in virulence-associated UPS and to define the composition of molecular machinery involved in release of confirmed and putative virulence factors, in this study we employed affinity purification and mass spectrometry coupled to microscopy-based subcellular localization and signal correlation quantification techniques to investigate protein complexes of eleven reported unconventionally-secreted putative and confirmed virulence factors, all predicted to be cytosolic. A subset of selected putative and confirmed virulence factors, along with their interaction partners, unequivocally associate to the surface of PV/PECs. Extended and validated interactomes point to a core PV/PECs-associated UPS machinery, which includes uncharacterized and Giardia-specific coiled-coil proteins and NEK kinases. Finally, a specific subset of the alpha-giardin protein family was invariably found enriched in all PV/PECs-associated protein interactomes, highlighting a previously unappreciated role for these proteins at PV/PECs and in UPS.

Taken together, our results provide the first characterization of a virulence-associated UPS protein complex in *Giardia lamblia* at PVs/PECs, suggesting a novel link between these primarily endocytic and feeding organelles and UPS at the parasite-host interface.

## Introduction

Unconventional protein secretion (UPS) is an umbrella term which defines secretion routes alternative to classic ER to Golgi secretion, guided by signal peptides (Balmer & Faso, 2021; Rabouille et al., 2012). To date, four UPS pathways have been formally described: protein secretion via self-sustained channel formation (UPS I), ABC transporter-mediated membrane passage (UPS II), vesicular export involving autophagy components (UPS III) and Golgi bypass of membrane proteins (UPS IV) (Balmer & Faso, 2021; Kim et al., 2018; Rabouille, 2017). UPS has also been reported in protist parasites and was investigated with respect to virulence in Leishmania (Denny et al., 2000; Maclean et al., 2012), Plasmodium (Cha et al., 2016; Ghosh et al., 2011) and Trichomonas (Miranda-Ozuna et al., 2016).

*Giardia lamblia* (*syn. Giardia intestinalis, Giardia duodenalis*) is a small-intestine protist parasite with a worldwide distribution. *Giardia* presents a simplified endomembrane system, with only five membrane-bound compartments, including an extensive ER, and no detectable Golgi apparatus (Elias et al., 2008; Hehl & Marti, 2004; Marti & Hehl, 2003) despite constitutive trafficking of secretory variant surface proteins (Faso & Hehl, 2011, 2019; McCaffery et al., 1994). The sole port of entry for fluid-phase material in the Giardia cell are the recently renamed (Santos et al., 2022) peripheral vacuoles/peripheral endocytic compartments (PVs/PECs), small (ca. 200nm) highly polymorphic organelles (Cernikova et al., 2018, 2019, 2020; Pipaliya et al., 2021; Zumthor et al., 2016). These organelles present a clearly endocytic molecular complement composed of clathrin assemblies, endocytic adaptor and lipid-binding proteins, ESCRT components and putative receptors (Cernikova et al., 2020a; Pipaliya et al., 2021; Zumthor et al., 2016b). PVs/PECs perform cycles of uptake and release of extracellular medium as they temporarily fuse with the plasma membrane (Cernikova et al., 2018; Zumthor et al., 2016b).

Virulence at the Giardia-host interface is poorly understood, with secreted cysteine proteases (CPs) and their documented negative impact on host cells (disruption of cell-cell adhesion, microbiota and the mucus layer, apoptosis of epithelial cells) as perhaps the best characterized example of virulence factors (Allain et al., 2019; Liu et al., 2018; G. Ortega-Pierres et al., 2018; M. G. Ortega-Pierres & Argüello-García, 2019). Recent reports document a large number of Giardia-derived proteins released in the extracellular space or found on the surface of Giardia cells (Davids et al., 2019; Dubourg et al., 2018; Ma’ayeh et al., 2017), in both axenic and co-cultivation conditions, with mostly unclear roles in virulence. However, the striking feature of these data is that, in contrast to CPs which present unequivocal sequences for secretion, the vast majority of proteins detected in secretome and surface proteome analyses of Giardia trophozoites are predicted to be soluble, non-secretory and intracellular, thus implicating UPS in their release. Among them are notable examples such as enolase (ENO) and arginine deiminase (ADI). ENO is perhaps one of the most studied candidate UPS substrate for its role as putative virulence factor in several parasitic species, including *Plasmodium, Trichomonas* and *Entamoeba* (Ahn et al., 2018; Ghosh et al., 2011; Miura et al., 2012; Tovy et al., 2010), although its role in pathogenesis is not fully understood (Pancholi, 2001; Ringqvist et al., 2008). In co-cultivation studies with intestinal epithelial cells and subsequent comparison to secretion in axenic cultures, Giardia-derived ENO and ADI were shown to be increased in the extracellular medium after incubation with host cells (Eckmann et al., 2000; Ma’ayeh et al., 2017; Ringqvist et al., 2008). In contrast to ENO, a more robust role for Giardia released ADI-mediated host arginine depletion has been defined, with a measurable negative impact on nitric oxide production, intestinal T-cell proliferation and dendritic cell cytokine secretion (Adam, 2021; Stadelmann et al., 2013).

Despite the fact that the Giardia-host interface is populated by many soluble non-secretory proteins, some of them with documented virulence function, no dedicated machinery nor mechanism(s) for UPS in Giardia are currently known. Speculations on the involvement of PV/PECs have been put forth previously (Benchimol & de Souza, 2022; Midlej et al., 2019; Moyano et al., 2019). We therefore hypothesized that, with PV/PECs at the host-pathogen interface and in direct communication with the extracellular environment, a dedicated UPS machinery could be found at PV/PECs for the release of unconventionally-secreted putative and confirmed virulence factors.

To test this hypothesis, we selected 11 putative UPS substrates with no detectable secretory signals and transmembrane domains, and characterized them in terms of subcellular localisation and protein interaction partners. We employed immunofluorescence assays, including co-labelling experiments and quantification of signal overlap, on Giardia trophozoites expressing epitope-tagged putative UPS substrate. This was followed by co-immunoprecipitation (co-IP) and mass spectrometry-based protein identification, to define and expand the UPS interactome. In line with the presented hypothesis, a subset of the selected putative virulence factors were found to localise to the surface of PVs/PECs in a tight interactome network with specific PV-PECs-associated NEK kinases and coiled-coil proteins. This network includes several alpha giardins, annexin homologs (Popa et al., 2018; Weiland et al., 2005) which also localise to PV/PECs. These data shed light on a novel link between unique endocytic compartments and virulence-associated UPS protein complexes at the parasite-host interface in *Giardia lamblia*.

## Results

### Selected putative UPS substrates localise to PV/PECs

As a first step towards testing the hypothesis that UPS in Giardia is linked to PV/PECs, we searched through reported datasets derived from secretome and surface proteome analyses (Davids et al., 2019; Dubourg et al., 2018; Ma’ayeh et al., 2017). We selected 11 putative UPS substrates (table 1) based on criteria extracted from GiardiaDB and other sources:

**Table 1.** Selected putative UPS substrates including histone H2A as a negative control for secretion and peroxiredoxin-1 as a predicted canonically secreted protein. Open reading frame (ORF) numbers and protein names are followed by the secretome or surface proteome study they were identified in (Davids et al., 2019; Dubourg et al., 2018; Ma’ayeh et al., 2017). The third column indicates if the protein is predicted to carry a signal peptide for secretion (SP) and/or a transmembrane domain (TMD), as retrieved from GiardiaDB. The last column extracts transcriptomics data from two RNA seq studies on *G. lamblia* WB-A (Franzén et al., 2013; Tolba et al., 2013) with transcript abundance expressed in in TPM (transcripts per million).

| ORF and protein name | found in secretome (sec.) or surface proteome (sur.) | SP or TMD | Unique transcript expression RNA seq WBA (Franzén et al.2013/ Tolba et al 2013) [TPM] |
| --- | --- | --- | --- |
| GL50803_11118 Enolase | sec. (Dubourg et al. 2017, Ma'ayeh et al. 2017) | no | 623.14/1019.83 |
| GL50803_112103 Arginine deiminase | sec. (Ma'ayeh et al. 2017) | no | 1100.54/1715.93 |
| GL50803_4812 beta giardin | sur. (Davids et al. 2019) and sec. (Dubourg et al. 2017, Ma'ayeh et al. 2017) | no | 2473.07/0 |
| GL50803_11654 alpha 1-giardin | sur. (Davids et al. 2019) and sec. (Dubourg et al. 2017, Ma'ayeh et al. 2017) | no | 783.76/2887.52 |
| GL50803_17153 alpha 11- giardin | sur. (Davids et al. 2019) and sec. (Dubourg et al. 2017, Ma'ayeh et al. 2017) | no | 398.57/1099.9 |
| GL50803_16453 carbamate kinase | sur. (Davids et al. 2019) and sec. (Ma'ayeh et al. 2017) | no | 1535.07/729.65 |
| GL50803_17327 Xaa-Pro dipeptidase | sec. (Ma'ayeh et al. 2017) | no | 212.2/39.26 |
| GL50803_3910 hypothetical | sec. (Ma'ayeh et al. 2017) | no | 157.87/236.14 |
| GL50803_112304 EF1-alpha | sec. (Dubourg et al. 2017, Ma'ayeh et al. 2017) | no | 161.61/76 |
| GL50803_12091 Macrophage migratic | sec. (Dubourg et al. 2017) | no | 140.06/16.53 |
| GL50803_6687 GAPDH | sec. (Ma'ayeh et al. 2017) | no | 258.27/982.49 |
| GL50803_27521 Histone H2A (negative control) | no | no | 340.53/461.96 |
| GL50803_15383 Peroxiredoxin-1-paralog (positive control) | sur. (Davids et al. 2019) and sec. (Ma'ayeh et al. 2017) | SP | 175.09/183.86 |

- Detection in the extracellular space (according to (Davids et al., 2019; Dubourg et al., 2018; Ma’ayeh et al., 2017))
- Absence of predicted signal peptide using prediction algorithms (SignalP using SP-HMM/SP-NN (Almagro Armenteros et al., 2019))
- Absence of predicted transmembrane domains (detected by TMHMM (Krogh et al., 2001))
- Availability of transcriptomics data (Franzén et al., 2013; Tolba et al., 2013).

Stably-transfected *G. lamblia* WB-A transgenic lines expressing epitope-tagged variants of all selected putative UPS substrates were generated, including transgenic lines expressing either epitope-tagged histone H2A or peroxiredoxin-1-paralog variants as negative or positive controls for canonical secretion, respectively. Analysis by immunofluorescence assays (IFA) followed by widefield microscopy of all transgenic lines show that, in contrast to epitope-tagged histone2A and peroxiredoxin-1-paralog which localise to the nuclei and to the ER respectively, and to beta-giardin which localises to the ventral disc, all other selected UPS protein substrates localise to the cytosol, as predicted (Figure 1). However, Xaa-Pro dipeptidase, alpha 1-giardin, alpha 11-giardin and enolase show additional deposition in close proximity to PV/PECs, as indicated by signal at the periphery of the cell and in the “bare zone” between nuclei (Chavez & Martinez-Palomo, 1995; Elmendorf et al., 2003; Friend, 1966) similar to *Gl*CHC, a previously characterized *Giardia* protein known to accumulate at PV/PECs (Cernikova et al., 2019, 2020a; Zumthor et al., 2016b). These proteins and their respective transgenic lines were selected for further investigation to better define subcellular localization of the corresponding reporter variants.

**Figure 1.**
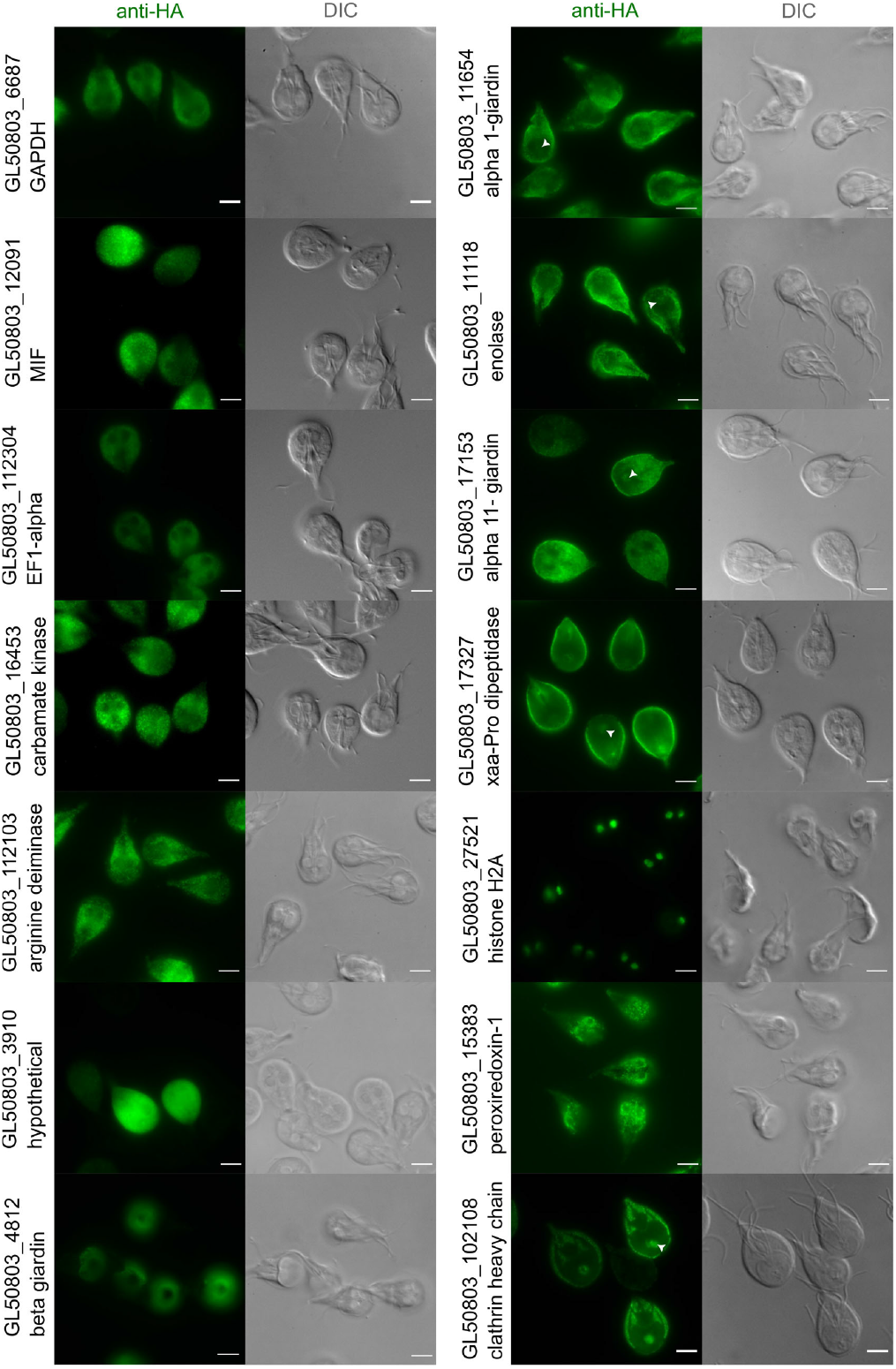
Selected putative UPS substrates localise to PV/PECs. Representative widefield light microscopy images of antibody-labelled HA epitope tagged UPS substrates and controls, expressed in Giardia trophozoites (anti-HA panels) including DIC images (Differential Interference Contrast). The first six transgenic lines on the left (GL50803_112103 Arginine deiminase, GL50803_12091 Macrophage migration inhibitory factor (MIF), GL50803_3910 hypothetical protein, GL50803_16453 carbamate kinase, GL50803_112304 TEF1 alpha, GL50803_6687 GAPDH (Glyceraldehyde 3-phosphate dehydrogenase)) show a cytosolic distribution, while GL50803_4812 beta-giardin shows localisation to the ventral disc. The first four transgenic lines on the right (GL50803_17327 Xaa-Pro dipeptidase, GL50803_11654 alpha 1-giardin, GL50803_17153 alpha 11-giardin, GL50803_11118 Enolase) show a proximity to PV/PECs. GL50803_27521 Histone H2A localises to nuclei while GL50803_15383 Peroxiredoxin-1 shows an ER localisation pattern. GL50803_102108 *Gl*CHC is included as a *bona fide* PV/PECs localised protein (Cernikova et al., 2020b; Santos et al., 2022; Zumthor et al., 2016a). The PV/PECs-enriched bare zone is highlighted with a white arrowhead. Scale bars: 5µm.

### Confocal microscopy and signal overlap analysis confirms UPS substrates at PV/PECs

As widefield microscopy analysis of transgenic antibody-labelled Giardia lines expressing tagged variants of Giardia proteins Xaa-Pro dipeptidase, alpha 1-giardin, alpha 11-giardin and enolase revealed localisation in close proximity to PV/PECs, we applied confocal microscopy analysis following immunofluorescence assays (IFA; Figure 2A). Giardia cells expressing Xaa-Pro dipeptidase, alpha 1-giardin, alpha 11-giardin and enolase present signal concentrated at the cell periphery and in the bare-zone between the nuclei.

**Figure 2.**
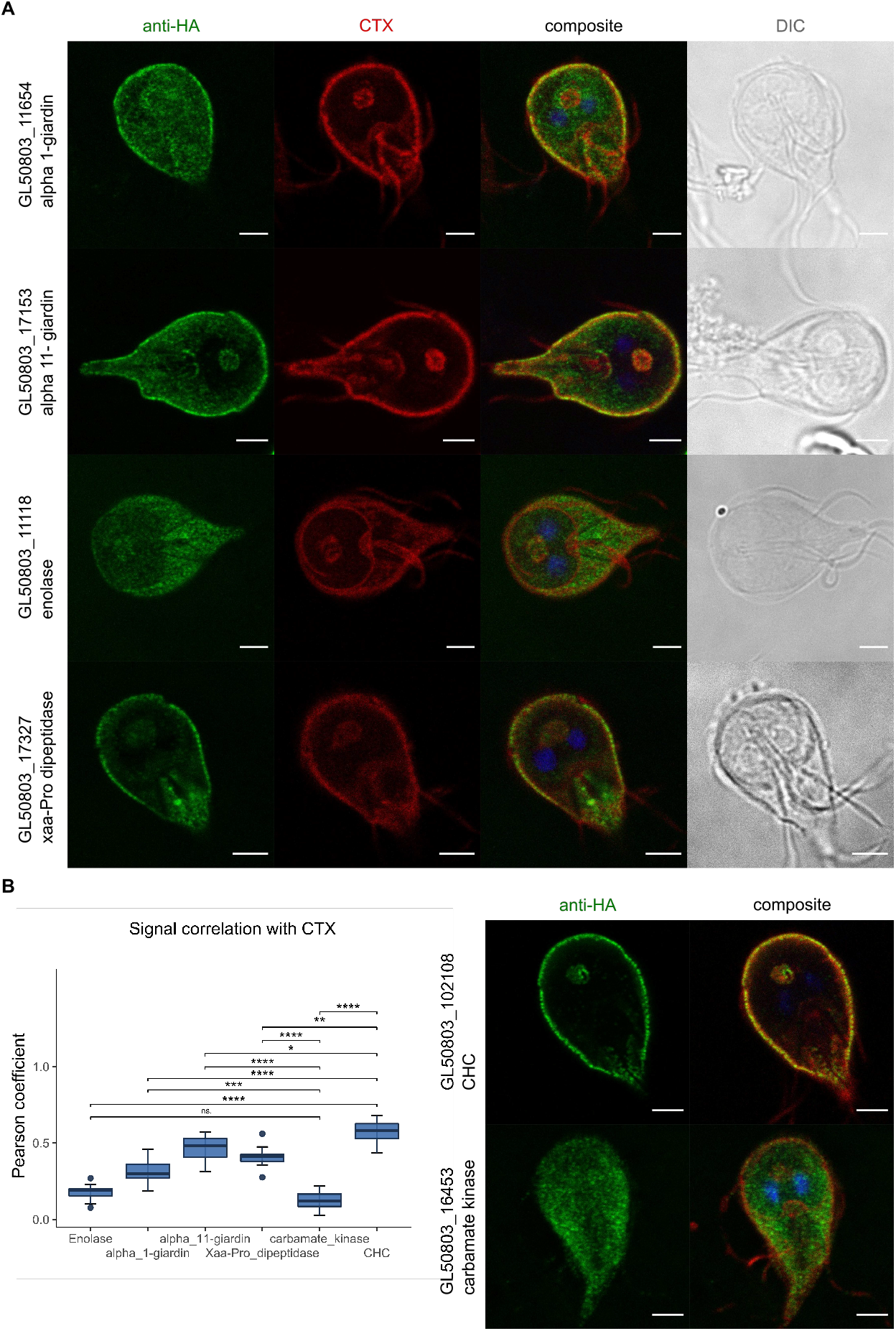
Confocal microscopy and signal overlap analysis confirms UPS substrates at PV/PECs. **(A)** Confocal microscopy images of antibody-labelled HA epitope tagged proteins expressed in transgenic Giardia trophozoites (anti-HA panels in green) including co-labelling with the PV/PECs membrane marker cholera toxin B, Alexa Fluor™ 594 Conjugate in red (CTX), a composite image of the two panels and the nuclei stained with DAPI and a DIC image. **(B)** Signal correlation for each transgenic line was quantified calculating the Pearson coefficient (N=10). A *t*-test was performed to compare all values to the cytosolic control carbamate kinase and GlCHC as a PV/PECs marker. Significance levels are indicated by asterisks. The data is displayed as a boxplot and n.s. indicates a non-significant difference (p-value > 0.05). The right panel shows representative single cell images of the control constructs. Scale bars: 3µm.

As a measure of the degree of association of reporter Xaa-Pro dipeptidase, alpha 1-giardin, alpha 11-giardin and enolase variants to PV/PECs, co-labelling experiments for the epitope-tagged variants with the plasma/PV membrane marker cholera toxin-b (CTX: (Corrêa et al., 2009; Zumthor et al., 2016b)) (Figure 2A) and subsequent signal correlation analysis using a dedicated macro script, was performed (Figure 2B and Supplementary data 2). Pearson coefficient values were calculated for each line (N=10) (Supplementary table 2), including control cell lines expressing epitope-tagged variants of *Gl*CHC and Giardia carbamate kinase (*Gl*CK), for a predominantly PV/PECs or cytosolic deposition pattern, respectively (Figure 2B, right panel). A moderate positive correlation is found for the tested lines, accounting for the substantial cytosolic protein pool for each reporter. Signal overlap analysis (Figure 2B, box plot) shows that enolase deposition presents the lowest correlation to CTX signal, with no significant deviation from the predominantly cytosolic *Gl*CK control line, despite clear PV/PECs association. However, signal overlap analysis for cells expressing Xaa-Pro dipeptidase, alpha 1-giardin and alpha 11-giardin variants indicates how these proteins differ significantly from *Gl*CK and approach *Gl*CHC as our reference marker for PV/PECs association.

**Table 2.** Top ten most abundant interactors of selected NEK kinases and coiled-coil proteins as measured by co-IP in limited cross-linking conditions. The top ten interactors of the investigated coiled-coil proteins and NEK kinases including the GL50803_6464 hypothetical protein which shows structural similarities to a coiled-coil protein, are listed. Interactors are sorted according to average relative iBAQ values. Highlighted in yellow is the affinity handle protein for each co-IP dataset and highlighted in blue are proteins detected in the interactome of PV/PECs-associated putative UPS substrates.GN: gene number.

| GL50803_16824 NEK Kinase |  |  | GL50803_15411 NEK Kinase |  |  |
| --- | --- | --- | --- | --- | --- |
| GN | Product Description | average %iBAQ | GN | Product Description | average %iBAQ |
| GL50803_16824 | NEK Kinase | 33.30 | GL50803_11654 | Alpha-1 giardin | 30.15 |
| GL50803_14521 | Peroxioredoxin 1 | 6.79 | GL50803_15411 | NEK Kinase | 22.50 |
| GL50803_112304/-12 | Elongation factor 1-alpha | 4.90 | GL50803_14551 | Alpha-6 giardin | 5.99 |
| GL50803_11654 | Alpha-1 giardin | 5.82 | GL50803_6687 | Glyceraldehyde 3-phosphate dehydrogenase | 3.20 |
| GL50803_114787 | Alpha-7.3 giardin | 2.29 | GL50803_15409 | NEK Kinase | 3.39 |
| GL50803_10311 | Ornithine carbamoyltransferase | 2.61 | GL50803_7796 | Alpha-2 giardin | 3.70 |
| GL50803_88765 | Cytosolic HSP70 | 1.98 | GL50803_14521 | Peroxioredoxin 1 | 2.94 |
| GL50803_93938 | Triosephosphate isomerase, cytosolic | 1.45 | GL50803_17153 | Alpha-11 giardin | 2.30 |
| GL50803_17153 | Alpha-11 giardin | 2.27 | GL50803_112304/-12 | Elongation factor 1-alpha | 2.48 |
| GL50803_6687 | Glyceraldehyde 3-phosphate dehydrogenase | 1.58 | GL50803_10311 | Ornithine carbamoyltransferase | 1.57 |
| GL50803_17570 | Elongation factor 2 | 1.15 | GL50803_114787 | Alpha-7.3 giardin | 1.26 |
| GL50803_11390 NEK Kinase |  |  | GL50803_15409 NEK Kinase |  |  |
| GN | Product Description | average %iBAQ | GN | Product Description | average %iBAQ |
| GL50803_17327 | Xaa-Pro dipeptidase | 29.27 | GL50803_15409 | NEK Kinase | 68.09 |
| GL50803_11390 | NEK Kinase | 17.72 | GL50803_11654 | Alpha-1 giardin | 3.04 |
| GL50803_17153 | Alpha-11 giardin | 9.00 | GL50803_17153 | Alpha-11 giardin | 4.17 |
| GL50803_113677 | Coiled-coil protein | 5.91 | GL50803_14521 | Peroxioredoxin 1 | 2.14 |
| GL50803_15409 | NEK Kinase | 6.57 | GL50803_10311 | Ornithine carbamoyltransferase | 1.95 |
| GL50803_11654 | Alpha-1 giardin | 3.77 | GL50803_137716 | Axoneme-associated protein GASP-180 | 1.43 |
| GL50803_11118 | Enolase | 4.77 | GL50803_112304/-12 | Elongation factor 1-alpha | 1.85 |
| GL50803_33769 | NADH oxidase lateral transfer candidate | 2.94 | GL50803_15411 | NEK Kinase | 1.59 |
| GL50803_10311 | Ornithine carbamoyltransferase | 3.18 | GL50803_16745 | Axoneme-associated protein GASP-180 | 1.03 |
| GL50803_6184 | Branched-chain amino acid aminotransferase lat | 3.93 | GL50803_17327 | Xaa-Pro dipeptidase | 1.04 |
| GL50803_14521 | Peroxioredoxin 1 | 1.48 | GL50803_114787 | Alpha-7.3 giardin | 0.96 |
| GL50803_17249 Coiled-coil protein |  |  | GL50803_6464 hypothetical protein (putative coiled-coil protein) |  |  |
| GN | Product Description | average %iBAQ | GN | Product Description | average %iBAQ |
| GL50803_15409 | NEK Kinase | 19.61 | GL50803_11654 | Alpha-1 giardin | 27.69 |
| GL50803_11654 | Alpha-1 giardin | 17.93 | GL50803_17153 | Alpha-11 giardin | 25.71 |
| GL50803_17249 | Coiled-coil protein | 16.94 | GL50803_6464 | hypothetical protein | 9.56 |
| GL50803_15411 | NEK Kinase | 11.69 | GL50803_16824 | NEK Kinase | 3.73 |
| GL50803_17153 | Alpha-11 giardin | 8.98 | GL50803_14521 | Peroxioredoxin 1 | 3.77 |
| GL50803_14551 | Alpha-6 giardin | 3.08 | GL50803_15409 | NEK Kinase | 3.21 |
| GL50803_112304/-12 | Elongation factor 1-alpha | 2.52 | GL50803_112304/-12 | Elongation factor 1-alpha | 2.62 |
| GL50803_14521 | Peroxioredoxin 1 | 2.26 | GL50803_10311 | Ornithine carbamoyltransferase | 2.05 |
| GL50803_10311 | Ornithine carbamoyltransferase | 1.75 | GL50803_7796 | Alpha-2 giardin | 1.87 |
| GL50803_7031 | Spindle pole protein, putative | 1.41 | GL50803_15411 | NEK Kinase | 2.07 |
| GL50803_7796 | Alpha-2 giardin | 1.18 | GL50803_17249 | Coiled-coil protein | 1.50 |
| GL50803_10167 Coiled-coil protein |  |  | GL50803_15591 Coiled-coil protein |  |  |
| GN | Product Description | average %iBAQ | GN | Product Description | average %iBAQ |
| GL50803_10167 | Coiled-coil protein | 52.82 | GL50803_15591 | Coiled-coil protein | 32.53 |
| GL50803_11654 | Alpha-1 giardin | 8.78 | GL50803_17153 | Alpha-11 giardin | 14.77 |
| GL50803_17153 | Alpha-11 giardin | 6.73 | GL50803_10311 | Ornithine carbamoyltransferase | 6.04 |
| GL50803_15411 | NEK Kinase | 3.71 | GL50803_112103 | Arginine deiminase | 3.89 |
| GL50803_15409 | NEK Kinase | 3.42 | GL50803_15409 | NEK Kinase | 2.62 |
| GL50803_14521 | Peroxioredoxin 1 | 3.33 | GL50803_8826 | Glucokinase | 2.41 |
| GL50803_11720 | hypothetical protein | 2.71 | GL50803_112304/-12 | Elongation factor 1-alpha | 1.66 |
| GL50803_112304/-12 | Elongation factor 1-alpha | 2.67 | GL50803_6687 | Glyceraldehyde 3-phosphate dehydrogenase | 1.42 |
| GL50803_11129 | hypothetical protein | 1.61 | GL50803_88765 | Cytosolic HSP70 | 0.94 |
| GL50803_10311 | Ornithine carbamoyltransferase | 1.87 | GL50803_14521 | Peroxioredoxin 1 | 0.82 |
| GL50803_14551 | Alpha-6 giardin | 1.55 | GL50803_8445 | NEK Kinase | 0.51 |

### UPS substrates at PVs/PECs participate in a core molecular complement

Having confirmed that *Giardia* putative UPS substrates Xaa-Pro dipeptidase, alpha 1-giardin, alpha 11-giardin and enolase are unequivocally and to varying degrees associated to the PV/PEC surface, we proceeded to define a UPS and PV/PECs-associated interactome. Epitope-tagged variants for each protein expressed and extracted from transgenic *Giardia* cells were used as affinity handles for co-immunoprecipitation (co-IP) in both limited cross-linking (to stabilize transient protein complexes) and native conditions, followed by mass spectrometry-based protein identification (Supplementary table 3). The ten most enriched interactors identified in each co-IP dataset, as measured by relative iBAQ (intensity-based absolute quantification) values across two independent biological replicates (Supplementary table 3), were used to build a UPS and PV/PECs-associated interactome network (Figure 3). Each selected PV/PECs-associated protein is connected to at least one of the other PVs/PECs-associated UPS substrates, with several shared interaction partners (Figure 3 and Supplementary tables 3 and 4). Alpha 1-giardin is the PVs/PECs UPS substrate with the most unique interactors, having tight connections to coiled-coil proteins, NEK kinases and alpha 6-giardin. Alpha 11-giardin showed reciprocal interactions to all three other selected PV/PECs-localised UPS substrates. Elongation factor 1-alpha and OCT were indiscriminately found across almost all UPS-related datasets in both types of co-IP conditions, indicating that abundance rather than specific interaction may account for their presence.

**Figure 3.**
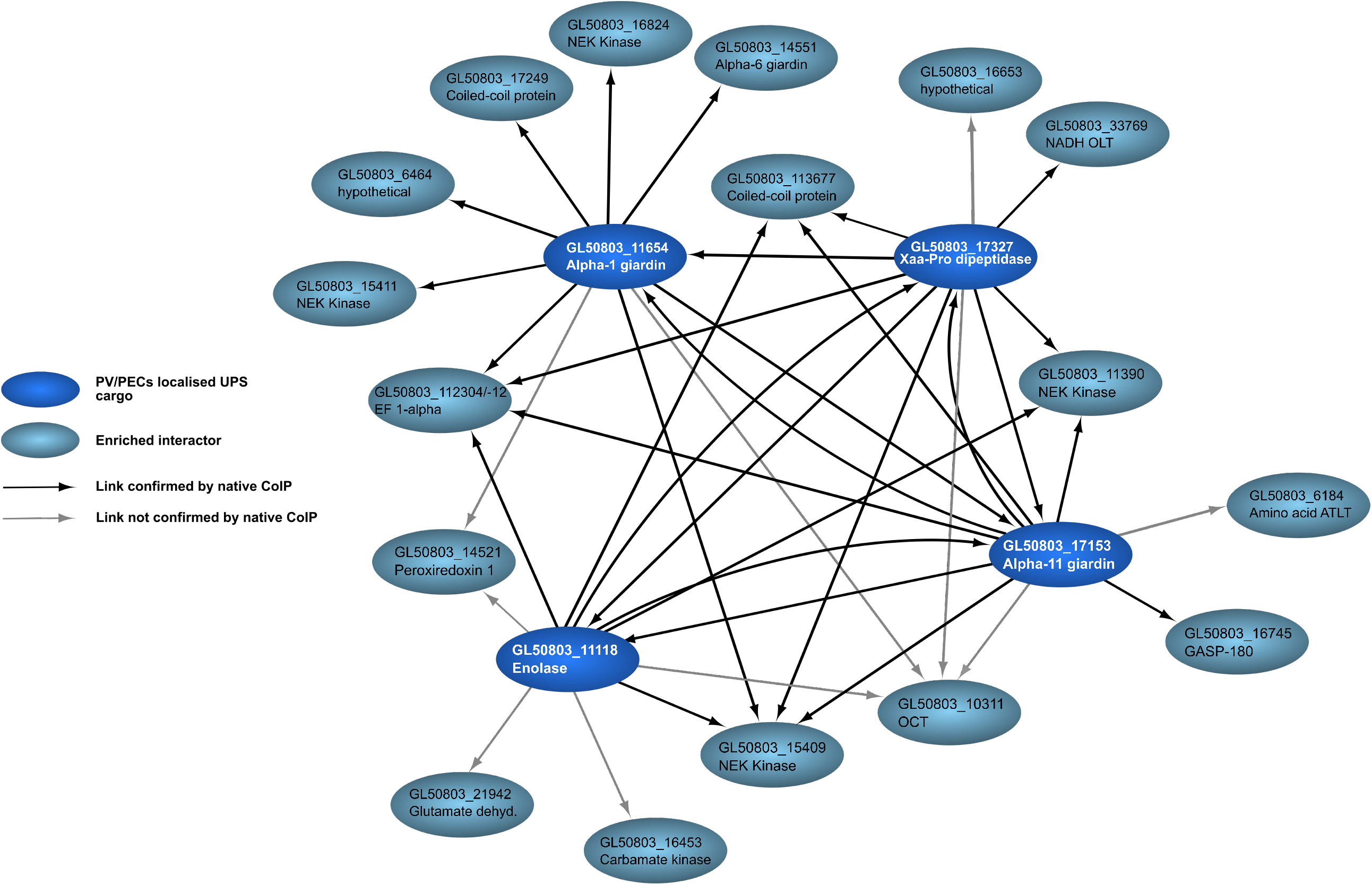
UPS substrates at PV/PECs participate in a core molecular complement. Depicted are the ten most enriched interactors in co-IP experiments for each PV-associated UPS substrate (in dark blue, GL50803_17327 Xaa-Pro dipeptidase, GL50803_11654 alpha 1-giardin, GL50803_17153 alpha 11-giardin, GL50803_11118 Enolase) based on relative iBAQ values. Directional interactions depicted in black were found in limited cross-linking conditions and confirmed by native co-IP; interactions in grey could not be confirmed by native co-IP.

Native co-IP confirmed 30 of the 40 displayed interaction links found by co-IP in limited crosslinking conditions. Data sets derived from co-IP of PV/PECs-associated UPS substrates proteins were compared to those for all selected UPS substrates, but beta-giardin, listed in table 1, and for ORF GL50803_27521 Histone H2A and ORF GL50803_15383 Peroxiredoxin-1-paralog as UPS controls, respectively (Supplementary table 3). Histone H2A is mainly associated to other histones (Supplementary table 3, pale blue) while the interactome of Peroxiredoxin-1 mainly consists of Protein disulphide isomerases (PDI) and other *bona fide* ER resident proteins (Supplementary table 3, pale green), as expected for a predicted canonically secreted protein carrying a secretory signal sequence. In stark contrast, NEK kinases and coiled-coil proteins are predominant in PV/PECs-associated data sets and absent from the list of top ten most enriched interactors for both UPS cytosolic substrates and UPS controls, in both types of co-IP setting (Supplementary table 3, pale yellow).

Given their documented roles as scaffolding proteins (NEK kinases) or tethering factors (coiled-coil proteins) (Gillingham & Munro, 2003; Manning et al., 2011), four NEK kinases (GL50803_15411, GL50803_16824, GL50803_15409, GL50803_11390) all predicted to be catalytically inactive (Manning et al., 2011), three annotated coiled-coil proteins (ORFs GL50803_10167, GL50803_15591, GL50803_17249) and one uncharacterized protein (GL50803_6464) were selected for further investigation. Homology prediction using the HHpred suite on ORF GL50803_6464 suggests this protein to also be a coiled-coil protein. Antibody labelling and widefield microscopy analysis of transgenic *Giardia* lines expressing epitope-tagged variants of the selected coiled-coil proteins and NEK kinases show reporter deposition at the cell periphery, with the exception of ORF GL50803_16824 NEK kinase, which appears to be predominantly cytosolic (Figure 4).

**Figure 4.**
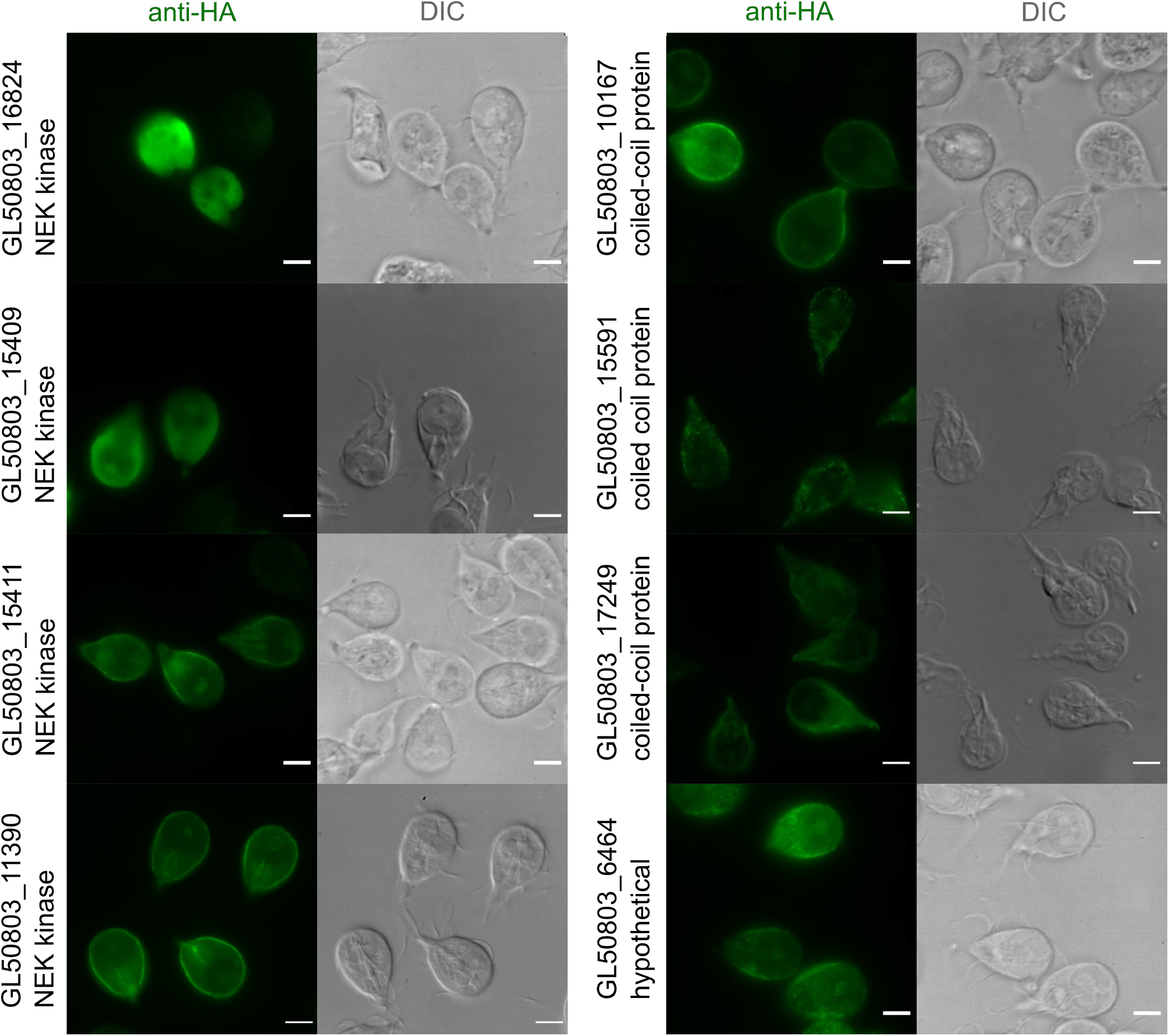
Selected coiled-coil proteins and NEK kinases localise to PV/PECs. Representative widefield light microscopy images of antibody-labelled HA epitope tagged proteins expressed in Giardia trophozoites (anti-HA panels) including DIC images. On the left, GL50803_16824 NEK kinase shows a cytosolic distribution while NEK kinases GL50803_15409, GL50803_15411 and GL50803_11390 show deposition in PV/PECs proximity. On the right the coiled-coil proteins, all showing PV/PECs proximity distribution patterns. GL50803_6464 hypothetical is considered a putative coiled-coil protein (CC) based on *in silico* prediction and shows a PV/PECs distribution pattern as well. Scale bars: 5µm.

### An expanded UPS interactome at PV/PECs is dominated by coiled coil proteins and NEK kinases as accessory proteins

Having determined a molecular complement for UPS substrates at PV/PECs, we sought to expand and validate this protein network by using epitope-tagged variants of the selected coiled coil proteins and NEK kinases as affinity handles for reverse or reciprocal co-IP experiments in cross-linking conditions. Table 2 lists the ten most enriched interaction partners for each affinity handle, as measured by relative iBAQ values (%iBAQ) across two independent biological replicates. Xaa-Pro dipeptidase, alpha 1-giardin, alpha 11-giardin and enolase are all identified and enriched across datasets derived from co-IP of the eight selected NEK kinases and coiled coil proteins, thus validating the initial UPS-associated protein interactome. Furthermore, three additional alpha giardins, namely GL50803_7796 alpha 2-giardin, GL50803_14551 alpha-6 giardin and GL50803_114787 alpha 7.3-giardin, were detected (Table 2).

Widefield microscopy analyses of labelled epitope-tagged variants of the selected coiled-coil proteins and NEK kinases (except ORF GL50803_16824 NEK kinase) show reporter deposition at the cell periphery and bare-zone. Similar to what was previously done for UPS substrates Xaa-Pro dipeptidase, alpha 1-giardin, alpha 11-giardin and enolase, we refined the subcellular localization analysis for all selected coiled-coil proteins and NEK kinases in co-labelling experiments with CTX, followed by confocal microscopy (Figure 5A) and signal overlap analysis using a dedicated macro script (Figure 5B and Supplementary data 2). We omitted ORF GL50803_16824 NEK kinase from this analysis due to its predominantly cytosolic deposition pattern.

**Figure 5.**
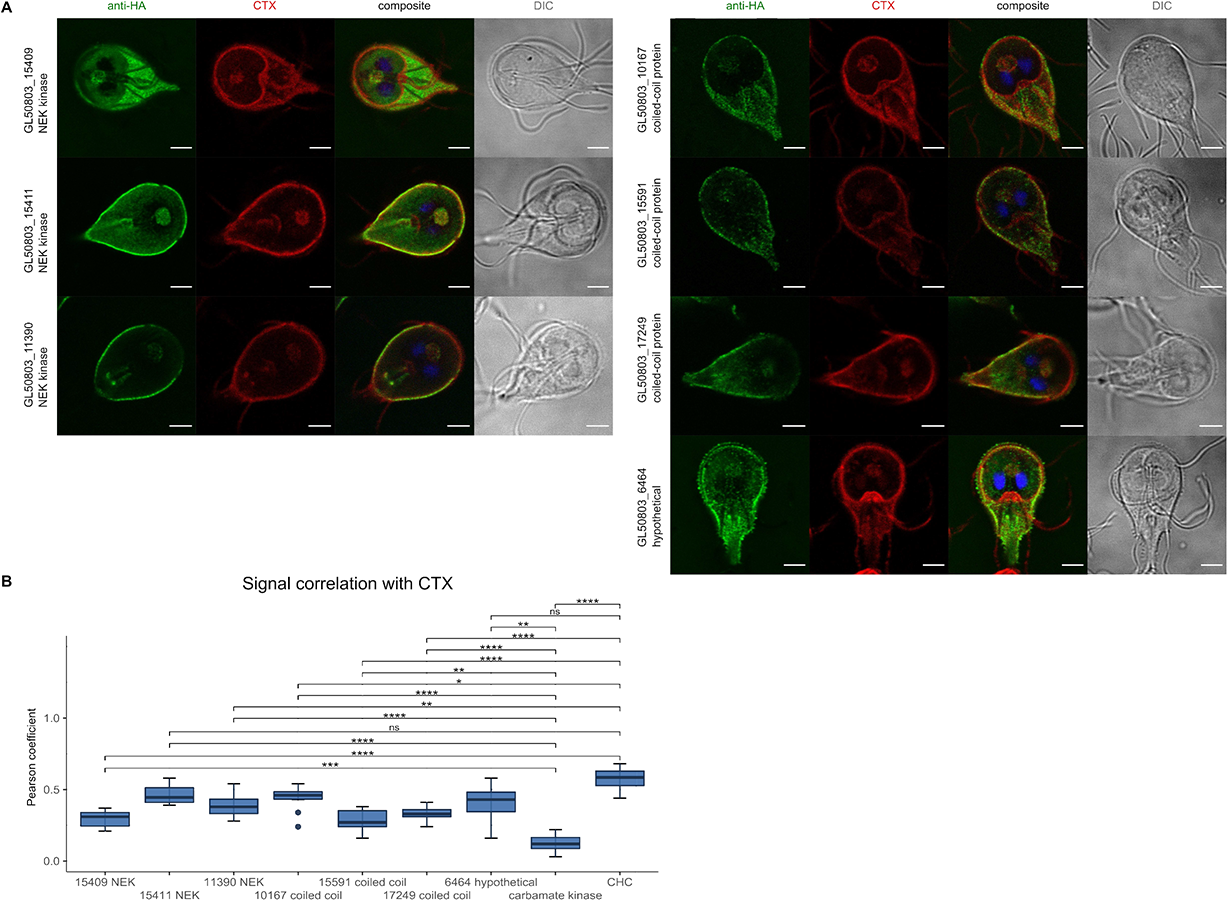
Confocal microscopy and signal overlap analysis confirms selected coiled-coils and NEK kinases at PV/PECs. (**A)** Confocal microscopy images of antibody-labelled HA epitope tagged proteins expressed in Giardia trophozoites (anti-HA panels in green) including co-labelling with the PV membrane marker cholera toxin B, Alexa Fluor™ 594 Conjugate in red (CTX), a composite image of the two panels and nuclei stained with DAPI and a DIC image. (**B)** Signal correlation was quantified calculating Pearson coefficient (N=10). A *t*-test was performed to compare all values to the cytosolic control carbamate kinase and *Gl*CHC. Significance levels are indicated by asterisks. The data is displayed as boxplots and n.s. indicates a non-significant difference (p-value > 0.05). Scale bars: 3µm.

All seven investigated proteins differ significantly from the cytosolic control carbamate kinase (Figure 2B, right panel) in their Pearsons coefficient values (Figure 5B and Supplementary table 2). GL50803_15411 NEK and GL50803_6464 hypothetical show high signal correlation, not significantly different from correlation values calculated for the *bona fide* PVs/PECs marker *Gl*CHC. Taken together, these and previous data validate the presence of a UPS interactome dominated by selected UPS substrates and associated to a defined set of coiled coil proteins and NEK kinases on the surface of PV/PECs.

## Discussion

### Identification of a UPS-associated protein complex at the PV/PECs surface

*Giardia* is known for its unique subcellular composition with only few endomembrane compartments (Acosta-Virgen et al., 2018; Benchimol & Souza, 2022; Faso & Hehl, 2019; Soltys et al., 1996; Tůmová et al., 2021). PV/PECs are positioned at the parasite-host interface as the sole gateway into the Giardia cell (Abodeely et al., 2009; Cernikova et al., 2018; Rivero et al., 2013; Zumthor et al., 2016; Santos et al., 2022) and with some speculation on their involvement in secretion or excretion (Benchimol & de Souza, 2022; Midlej et al., 2019; Moyano et al., 2019).

In this report, the hypothesis that, although primarily endocytic, PV/PECs may also be involved in release of putative and confirmed virulence factors which are not natively engineered for canonical i.e. ER-mediated secretion, was formulated. In line with this hypothesis, microscopy analyses show that at least four previously reported putative UPS substrates namely, Xaa-Pro dipeptidase, alpha 1-giardin, alpha 11-giardin and enolase (Davids et al., 2019; Dubourg et al., 2018; Ma’ayeh et al., 2017), are associated to the surface of the PV/PECs organelle system while maintaining a sizable cytosolic pool, as reflected in the signal correlation analysis using CTX as a marker for PV/PECs membranes. Furthermore, validated co-IP data was used to build a PV/PECs-associated UPS interactome which highlights the strong degree of association amongst the selected PV/PECs-associated UPS substrates themselves as well as interaction with a defined set of coiled-coil proteins and NEK kinases, in both limited-crosslinking and native co-IP experiments. NEK kinases likely play an important role in *Giardia* as they constitute a large fraction of the predicted Giardia kineome (Manning et al., 2011). Although predicted to be catalytically inactive, the NEK kinases found in the PV/PECs-associated UPS interactomes may have scaffolding functions (Manning et al., 2011) collaborating with coiled-coil proteins as tethers (Sztul et al., 2006), either within the complex or to maintain the complex at the PV/PECs membrane (Figure 6). Further investigations will be needed to determine the exact role for these protein families at PV/PECs in connection to UPS.

**Figure 6.**
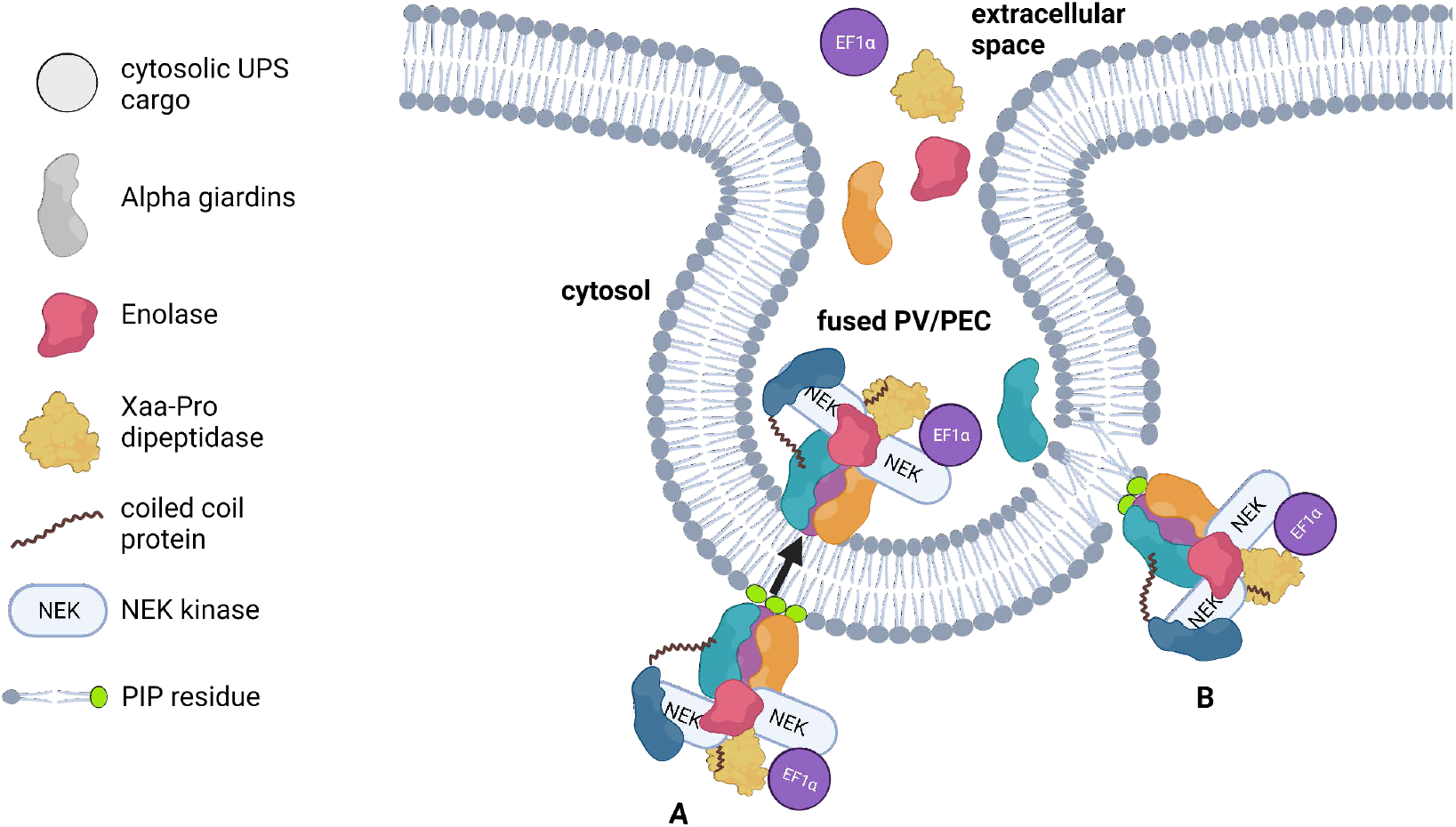
UPS, PV/PECs and alpha-giardins: a testable working model. This working model proposes alpha-giardins as both substrates and mediators of PV/PECs-associated UPS and is based on results presented in this and previous studies on annexins. In this model, alpha giardins are bound to the PV/PEC membrane, possibly via PIPs (green residues) (Goebeler et al., 2006; Hayes et al., 2004; Weingärtner et al., 2012) reported at the PV/PECs surface (Cernikova et al., 2020b), thus anchoring a UPS-related complex to the PV/PECs membrane. **(A)** In one scenario, alpha-giardins and associated UPS substrates are secreted as a protein complex, requiring the intervention of scramblases and/or flippases to invert topology, allowing for release in the PV/PECs lumen. **(B)** Alternatively, alpha giardin-mediated membrane destabilization would facilitate release of individual PV/PECs-associated UPS substrates to organelle lumina. In both scenarios **(A)** and **(B)**, release to the extracellular space would occur upon fusion of PV/PECs with the PM which is known to occur asynchronously and cyclically (Gaechter et al., 2008). EF1α: elongation factor 1α. This figure was created with Biorender.com

### A hypothesis on the role for alpha-giardins in PV/PECs membrane traversal

The co-IP datasets do not report on significant enrichment for transporters and/or components of vesicular carriers, which could have provided clues on how traversal of the PV/PEC membrane and extracellular release of Xaa-Pro dipeptidase, alpha 1-giardin, alpha 11-giardin and enolase is achieved. On the other hand, a striking feature of the co-IP datasets resides in the apparent enrichment for alpha-giardins as part of the UPS complex at PVs/PECs, including putative UPS substrates alpha 1-giardin and alpha 11-giardin which interact reciprocally and are both indirectly associated to alpha-giardins 2, 6 and 7.3.

Alpha-giardins are Giardia annexin orthologs (Morgan & Fernández, 1995). Following calcium-dependent binding of negatively-charged (acidic) phospholipids (Weeratunga et al., 2012), annexins, usually organized in oligomers, can destabilize biological membranes and even insert into them, inducing leakage in liposome-based experiments (Gerke & Moss, 2002; Luecke et al., 1995; Popa et al., 2018). In a recently-published review, mammalian annexins are discussed in the context of UPS due to their extracellular release without any detectable canonical secretory signal (Popa et al., 2018). Taking these previous reports on annexins and alpha-giardins into account, and combining them with data presented in this report, a working model is proposed for alpha-giardins as both substrates and mediators of PV/PECs-associated UPS (Figure 6).

The model allows for at least two testable predictions. The first is that alpha-giardins directly associate to PV/PEC membranes, without the need for membrane adaptors and in contrast to what was previously shown for clathrin assemblies (Zumthor et al., 2016b). Support for this prediction comes from previous data showing how some annexins (Goebeler et al., 2006a; Hayes, Merrifield, Shao, Ayala-Sanmartin, D’Souza Schorey, et al., 2004; Weingärtner, Kemmer, Müller, Zampieri, Gonzaga dos Santos, et al., 2012) bind phosphorylated derivatives of the minor membrane phospholipid phosphatidyl-inositols (PIPs) (Ridgway, 2016). PIPs were already detected at the PV/PECs surface (Cernikova et al., 2019, 2020a). Investigating phospholipid binding affinity of PV/PECs-associated alpha-giardins would begin to address the question of their role at this compartment and in the context of UPS. An essential role for PI(4,5)P_2_ has already been shown in the unconventional and self-sustained secretion of FGF-2 (UPS pathway I) (la Venuta et al., 2015; Legrand et al., 2020; Steringer et al., 2015; Wegehingel et al., 2008).

The second testable prediction is that alpha-giardins and associated UPS substrates are secreted as a protein complex (Figure 6A), similar to secretion of annexin 2 clusters bound to the PM by scramblase-mediated lipid rearrangement (Stewart et al., 2018). Native co-IP of *extracellular* alpha-giardins 1 and 11 would allow to determine whether PV/PECs-associated UPS substrates are released together, as also suggested by the native co-IP data presented in this report, or whether alpha giardin-mediated membrane destabilization facilitates leakage of UPS substrates into the PV/PEC lumen for individual release (Figure 6B). *In vitro* liposome-based testing platforms may lend support to one or the other scenario. Taken together, PV/PEC-associated alpha-giardins are proposed to play a role in membrane traversal of selected UPS substrates. Further investigations will be required to determine whether this constitutes a novel and perhaps unique UPS pathway.

## Materials and methods

### Giardia culturing conditions and transfection

*Giardia lamblia* trophozoites of strain WBA C6 (ATCC 50803) were axenically cultured according to previously established protocols (Cernikova et al., 2020a; Morf et al., 2010; Pipaliya et al., 2021; Zumthor et al., 2016b). Cells were grown in standard *Giardia* growth medium at 37°C and passaged every two to three days when cultures had reached confluency. Episomal or stable transfection of wild-type parasites with circular or linearized plasmid vectors (pPacV-Integ-based) was performed using electroporation at 350V, 960 µF, 800Ω followed by selection with puromycin for one week at 50ug per ml (InvivoGen). Expression of transfected constructs was controlled using immunofluorescence microscopy assays (IFA) and immuno-blotting.

### Construct synthesis

General information on ORFs including transmembrane domain and signal peptide predictions were gathered from the respective gene pages on GiardiaDB (table 1 and supplementary table 4). ORFs (Supplementary table 1) were amplified via PCR on genomic DNA from *Giardia* lamblia trophozoites of strain WBA C6 (ATCC 50803) and tagged at the C-terminus with a hemagglutinin (HA) - epitope tag using dedicated oligonucleotide pairs (Supplementary table 1). Restriction sites *XbaI* and *PacI* were introduced using the aforementioned oligonucleotides. Resulting amplicons for all constructs were cloned into a modified pPacV-Integ (Štefanić et al., 2009) vector (supplementary material). All proteins were expressed under their putative endogenous promoters defined as a fragment 200 to 250 bps upstream of the ORFs’ start codon. There are two gene copies of Elongation factor 1-alpha present in *Giardia* which differ only by their up- and downstream regions (GL50803_112304 and GL50803_112312) while thier coding regions differ only by two nucleotides, which do not affect the predicted protein sequence (Skarin et al 2011). As these two genes give rise to the same protein we decided to only further investigate GL50803_112304. The two proteins could not be distinguished by the MS analysis. For this reason, we indicate Elongation factor 1-alpha with the number GL50803_112304/-12 in these data sets.

### Immunofluorescence assays and microscopy analysis

Immunofluorescence assays were performed as described in previous studies (Cernikova et al., 2020a; Morf et al., 2010; Pipaliya et al., 2021; Zumthor et al., 2016b). Briefly, transgenic lines were grown to confluency in 12 ml Nunc polystyrene culture tubes (Thermo Fisher Scientific) and then cooled on ice to detach for 30-60 min. Tubes were hit on a soft surface to detach all cells and then centrifuged at 900*g* for 10 minutes. The cell pellet was washed in PBS (phosphate-buffered saline) and transferred to 1.5 ml Eppendorf tubes where the cells were fixed for a minimum of one hour or overnight in 3% formaldehyde (Sigma) in PBS. Alternatively, before fixation, cell pellets were resuspended in 40 µl culture medium and incubated with 8 µl of 1 µg/ml cholera toxin B, Alexa Fluor™ 594 Conjugate (Cat. No C22842, Thermo Fisher) at 37°C for 1h, washed in PBS and then fixed. After fixation, samples were quenched in 0.1M Glycine in PBS for 5 minutes before permeabilization in 1 ml of 2% BSA (bovine serum albumin) +0.2% Triton-X-100 in PBS, for 20 minutes at room temperature. Cells were then incubated with antibodies in 2% BSA+0.1%Triton-X-100 in PBS. Primary antibody: rat-derived monoclonal anti-HA antibody (dilution 1:250; Roche). Secondary antibody: goat-derived anti-Rat IgG (H+L) conjugated to Alexa Fluor 488 (AF488) (dilution 1:250; Thermo Fisher). Samples were incubated at room temperature for 1h in the dark. After each antibody incubation, samples were washed twice in 1% BSA+0.05% TX-100 in PBS. Cells were then carefully resuspended in *ca*. 30 µl Vectashield (Reactolab) containing 4′-6-diamidino-2-phenylindole (DAPI) as a nuclear DNA label. Cells were imaged at a Leica DM5500 widefield microscope and a Leica SP8 confocal microscope configured with white light lasers, general at full cell diameter.

Signal correlation analysis using Pearson’s coefficients was performed with Fiji ImageJ for 10 cells per transgenic line on a ROI defined in a standardised fashion by the cell contour (Fiji macro in supplementary data 2; Schindelin et al. 2019). Statistical analysis (t-test) was performed in R and visualisation in form of boxplots was done in R and Inkscape vector graphics editor (R Core Team, 2021). Pearson values were compared to values from a cytosolic control (carbamate kinase) and a construct previously shown to localise PV/PECs (clathrin heavy chain *Gl*CHC). Significance levels are indicated by asterisks (* P-value <=0.05, ** P-value <= 0.01, *** P-value <=0.001, **** P-value <=0.0001). “n.s.” indicates a non-significant difference (p-value > 0.05).

### Co-immunoprecipitation with limited cross-linking and in native conditions

Protocols for co-immunopreciptation (co-IP) under limited cross-linking conditions as well as native conditions were adapted from previous studies (Cernikova et al., 2020a; Zumthor et al., 2016b). *Giardia* trophozoites expressing tagged reporter constructs (minimal two independent biological replicates per line) as well as wild-type control WBA cells were grown in one T-25 flask per line, harvested and resuspended in PBS to reach a final volume of 10ml. For protein cross linking, cells were pelleted and resuspended in 2.25% formaldehyde in PBS and incubated for 30min at room temperature on a rotating shaker. Cells were then washed in PBS and incubated for 15min with 10ml PBS + glycine 100mM for quenching. Cells were then resuspended in 5 ml RIPA-SDS buffer (50mM Tris (pH 7.4), 150mM NaCl, 1% IGEPAL, 0.5% sodium deoxycholate, 0.1% SDS, and 10mM EDTA. 100μL) with added 0.1M phenylmethylsulphonyl fluoride (PMSF) and 50 µl protease inhibitor solution (Sigma). For controlled cell rupture, the resulting cell mixture was sonicated twice for 30 sec (60 pulses, 4 output control, 40% duty cycle) in the cold. After sonication the samples were incubated at 4°C on a rotating shaker for *ca*. 2h and then centrifuged and filtered (0.2μm WWPTFE membrane). To the filtrate 5ml RIPA-Triton solution (50mM Tris pH7.4, 150mM NaCl, 1% IGEPAL, 0.5% Na deoxycholate, 1% Triton x100, 10mM EDTA) and 40ul anti-HA agarose bead slurry (Thermo Fisher) were added and incubated overnight at 4°C on a rotating shaker. The beads were washed three times in TBS + 0.1% Triton X-100 and three times in PBS. Beads were stored dry at -20°C.

For the co-immunoprecipitation in native conditions, four transgenic UPS cargo lines showing a PV/PEC localization (GL50803_11118 Enolase, GL50803_11654 alpha 1-giardin, GL50803_17153 alpha 11-giardin, GL50803_17327 Xaa-Pro dipeptidase), a cytosolic control (GL50803_16453 carbamate kinase) and wild-type control WBA were grown and harvested as described above. The cell pellets were resuspended in 5ml PBS supplemented with 100µl PMSF 0.1M + 50µl protease inhibitor solution before the samples were sonicated, as described above. After sonication 250 µl 20% Triton X-100 was added and the samples were incubated at 4°C on a rotating shaker for *ca*. 2h. The samples were then centrifuged and filtered (0.2μm WWPTFE membrane) before the addition of 40µl anti-HA agarose bead slurry (Thermo Fisher) and incubation overnight. All samples were then subject to mass spectrometry-based protein identification.

### Liquid Chromatography Mass Spectrometry (LC/MS) and co-IP data analysis

Mass spectrometry and protein identification was performed by the Core Facility for Proteomics & Mass Spectrometry of the University of Bern. In a first step the samples were resuspended in 8M Urea with 50mM Tris-HCl at pH 8 and then reduced at 37°C with DTT 0.1M with 100mM Tris-HCl at pH 8 and alkylated at 37°C in the dark with IAA 0.5M and 100mM Tris-HCl for 30 mins. The slurry was then diluted four times with 20mM Tris-HCl with 2mM CaCl_2_ before digestion overnight with 100 ng sequencing grade trypsin (Promega). The samples were then centrifuged for peptide extraction from the supernatant which were then subject to liquid chromatography LC-MS (PROXEON coupled to a QExactive mass spectrometer, Thermo Fisher Scientific). µPrecolumn C18 PepMap100 (5μm, 100 Å, 300 μm×5mm, Thermo Fisher Scientific, Reinach, Switzerland) was used to trap the peptides and then they were separated by backflush on a C18 column (5 μm, 100 Å, 75 μm×15 cm, C18) by applying a 40-min gradient of 5% acetonitrile to 40% in water, 0.1% formic acid, at a flow rate of 350 nl/min. Full Scan was set at a resolution of 70 000, an automatic gain control (AGC) target of 1E06, and a maximum ion injection time of 50 ms. The following settings were applied with the data-dependent method for precursor ion fragmentation: resolution 17,500, AGC of 1E05, maximum ion time of 110 ms, mass isolation window 2 m/z, collision energy 27, under fill ratio 1%, charge exclusion of unassigned and 1+ ions, and peptide match preferred, respectively. MaxQuant (v. 1.6.14.0) was used for MS data interpretation against a *Giardia lamblia* database (Giardiadb v. 47) using the default MaxQuant settings. Transgenic lines were compared to wild type samples to be aware of possible contaminants. MS hits were sorted by their abundance according to the intensity-based absolute quantification (iBAQ) values. Relative abundance was then calculated from the total iBAQ for each protein hit (%iBAQ= iBAQ/ΣiBAQ*100). To validate the hits, replicates from each line were intersected and only hits found in both/all the data sets in similar relative abundance were further analysed. The average relative iBAQ was calculated by adding the relative iBAQ values for each hit and dividing them by the number of replicates. Average relative iBAQ was then sorted from highest to lowest. The ten most enriched hits aside from the bait protein were visualized with Cytoscape (Shannon et al. 2003). Proteomics data are deposited to the ProteomeXchange Consortium via the PRIDE (Perez-Riverol et al., 2022) partner repository with the dataset identifiers **PXD035195** (cross-linking co-IP conditions) and **PXD035190** (native co-IP conditions) (supplementary table 5). Data in both PXD035195 and PXD035190 can be accessed through http://www.ebi.ac.uk/pride by entering PXD035195 reviewer account details (username, password: o8PUqwbH) and PXD035190 reviewer account details (username, password: J1Q36yqx).

## Supplementary information

**Supplementary table 1**. List of oligonucleotide pairs used for ORF amplification

**Supplementary table 2**. Pearson coefficients calculated in image analysis

**Supplementary table 3**. Top 10 most enriched substrates in Co-IP mass spectrometry experiments. Tab 1: results of Co-IP/MS under crosslinking conditions with NEK kinases and coiled-coil proteins highlighted in pale yellow. In the histone negative control other histones are highlighted in pale blue. In Peroxiredoxin-1 (positive control for canonical secretion), *bona fide* ER resident proteins are highlighted in pale green. Tab 2: results of Co-IP/MS under native conditions with NEK kinases and coiled-coil proteins highlighted in pale yellow. GN: gene number.

**Supplementary table 4**. Information of highest enriched interactors in interactome network.

**Supplementary table 5**. Specifications on samples used for MS analysis of co-IP experiments deposited on PRIDE with identifiers PXD035195 and PXD035190.

**Supplementary data 1**. Construct sequences in .ape format, accessible with the open-source software [APE](https://jorgensen.biology.utah.edu/wayned/ape/) (Davis & Jorgensen, 2022).

**Supplementary data 2**. FIJI Macro for signal correlation analysis on ROI.

## Author contributions

EAB, CDW and CF designed experiments. EAB and CDW performed all experiments. EAB produced the figures and tables, did the data analysis, acquired all microscopy data and wrote the first draft of this manuscript. CF and EAB revised the manuscript. All authors authorize this submission.

## Acknowledgments

We thank the Microscopy Imaging Center of the University of Bern, particularly Dr. Yury Belyaev for training and advice, and members of BIOP of the EPFL and ScopeM of ETH Zurich for their help setting up an image analysis pipeline during the ZIDAS workshop. We thank the Proteomics and Mass Spectrometry Core Facility (PMSCF) of the Department for Biomedical Research at the University of Bern for generating all proteomics data. Funding for this project is provided by Swiss National Science Foundation grant numbers PR00P3_179813, PR00P3_179813/2 and PR00P3_179813/3 awarded to CF.

