## Supplementary data and tables for "A core UPS molecular complement implicates unique endocytic compartments at the parasite-host interface in *Giardia lamblia*": Supplementary data 2_FIJI Macro for signal correlation analysis on ROI.ijm

```
//define channels
DAPI=1
marker=2
HA_Ch=3

//choose image
imagetitle= getTitle();
print(imagetitle);


//define a ROI along cell membrane
selectWindow(imagetitle);
run("Duplicate...", "duplicate channels=2");
run("Make Binary", "method=Mean background=Dark only black");
run("Dilate");

waitForUser("check mask")
run("Analyze Particles...", "size=20-Infinity show=Outlines display add slice");
if (roiManager("count")>0){
	roiManager("select", 0);
	run("Enlarge...", "enlarge=0.2");
	roiManager("Update");
	print("ROIselected");

	//calculate pearsons correlation coefficient on marker channel and HA channel on whole image
	selectWindow(imagetitle);
	run("BIOP JACoP", "channel_a="+marker+" channel_b="+HA_Ch+" threshold_for_channel_a=Otsu threshold_for_channel_b=Otsu manual_threshold_a=0 manual_threshold_b=0 get_pearsons get_fluorogram costes_block_size=5 costes_number_of_shuffling=100");
	print("pearsonresult1");
	
	//run costes analysis on ROI for channel 2 (marker) and 3 (HA)
	selectWindow(imagetitle);
	run("BIOP JACoP", "channel_a="+marker+" channel_b="+HA_Ch+" threshold_for_channel_a=Otsu threshold_for_channel_b=Otsu manual_threshold_a=0 manual_threshold_b=0 perform_costes_randomization perform_costes_randomization_mask show_costes_plot show_costes_random_image costes_block_size=5 costes_number_of_shuffling=100");


}
```
